# Emergence of *mutH*^V76G^ among longitudinal carbapenem resistant *Klebsiella pneumoniae* causing long-term colonization and recurrent infection disrupts DNA mismatch repair and results in a hypermutator phenotype

**DOI:** 10.1101/2025.11.17.688822

**Authors:** Shaoji Cheng, Cornelius J Clancy, Giuseppe Fleres, Hassan Badrane, Matthew J Culyba, Anthony Newbrough, Liang Chen, M. Hong Nguyen

**Author notes:** Corresponding author: M. Hong Nguyen, M.D., Biomedical Science Tower #1 (Starzl), 200 Lothrop Street, Suite E-1051, Pittsburgh, PA 15213.

## Abstract

Although hypermutation due to Mut protein mutations that disrupt DNA mismatch repair has been characterized in some bacteria, its mechanisms and consequences in *Klebsiella pneumoniae* remain poorly defined. We analyzed 11 longitudinal KPC-3 carbapenemase-producing, ST258 *K. pneumoniae* isolates collected over ∼4 years from an immunocompromised patient with chronic colonization and recurrent infections. After ∼3.3 years, isolates developed ceftazidime–avibactam (CZA)-resistance with restored carbapenem susceptibility, coinciding with emergence of a V76G substitution in a highly-conserved motif in the core of MutH endonuclease. Compared with earlier isolates, *mutH*^V76G^-carrying isolates showed greater within-host genomic diversification (69-179 *vs.* 2–12 SNP differences) and acquired *bla*_KPC-3L169P_, encoding an KPC Ω-loop substitution that mediates CZA resistance and re-establishes carbapenem susceptibility. *mutH*^V76G^ isolates exhibited stepwise increases in meropenem-vaborbactam (MVB) and cefiderocol minimum inhibitory concentrations, plausibly linked to substitutions in KPC, OmpK36 porin, CirA iron transporter and/or EnvZ kinase. Clinical *mutH*^V76G^ isolates and CRISPR-engineered *mutH*^V76G^ mutants were hypermutators based on rifampin mutational frequencies. Using isogenic mutant and parent strains, we confirmed that *mutH*^V76G^ accelerated evolution of CZA and MVB resistance *in vitro* and *in vivo*, promoted transfer and uptake of resistance plasmids, and improved bacterial fitness during mouse infections. Resistance evolution in mice recapitulated clinical trajectories, with *bla*_KPC-3_ and *ompK36* mutations emerging under CZA and MVB exposure, respectively. Phenotypes of *mutH*^V76G^ and *mutH*-null strains were comparable, indicating that the V76G substitution largely abrogates MutH function. Our findings reveal MutH-mediated hypermutation as an adaptive mechanism in *K. pneumoniae*, enabling rapid antibiotic resistance development and plasmid acquisition without fitness cost.

**Importance:** Hypermutator bacteria pose a formidable clinical threat by rapidly evolving antibiotic resistance and adapting within the human host. *Klebsiella pneumoniae* is a major cause of multidrug-resistant infections, yet the contribution of hypermutation to its evolution remains poorly characterized. Analyzing *K. pneumoniae* isolates collected over ∼4 years from a chronically infected/colonized patient, we demonstrate that emergence of a mutation in *mutH* (*mutH*^V76G^), a DNA mismatch repair gene, results in hypermutation phenotypes and rapid accumulation of gene mutations. Both clinical and lab-engineered *mutH*^V76G^ mutant strains rapidly acquire resistance or reduced susceptibility to new antibiotics like ceftazidime-avibactam, meropenem-vaborbactam and cefiderocol, due to mutations in carbapenemase (*bla*_KPC-3_), porin (*ompK36*) and other genes. *mutH*^V76G^-driven hypermutation also enhances horizontal transfer of resistance plasmids and improves *K. pneumoniae* fitness during mouse infections. This study is important for understanding *K. pneumoniae* hypermutation as a potent mediator of antibiotic resistance and other phenotypes relevant to human infections.

## Introduction

Carbapenem-resistant Enterobacterales (CRE) present an urgent global health threat. *Klebsiella pneumoniae* is the most common CRE in the United States. Carbapenem resistance in *K. pneumoniae* is most often mediated by *K. pneumoniae* carbapenemases (KPCs) and other carbapenemases. Over the past decade, novel β-lactam–β-lactamase inhibitor combinations, such as ceftazidime–avibactam (CZA) and meropenem–vaborbactam (MVB), have reduced mortality in serious CRE infections.^1–4^ However, resistance to these agents emerges in ∼5-8% of treated patients. CZA and MVB resistance most often arises from mutations within the KPC Ω-loop and OmpK36 porin, respectively.^2,5^ Chronic gastrointestinal (GI) colonization and recurrent invasive infections, typically originating from the GI reservoir, are common among survivors of CRE infections.^6–8^ There is a critical need to better understand mechanisms driving within-host resistance evolution and adaptation of CR-*K. pneumoniae* (CRKP) and other CRE.

In natural bacterial populations, ∼1-5% of strains may exhibit high spontaneous mutation rates.^9^ The frequency of hypermutation strains is higher in certain settings, including during chronic infections by bacteria like *Pseudomonas aeruginosa* and *Staphylococcus aureus*.^10–12^ The hypermutation phenotype is most often caused by mutational inactivation of methyl-directed DNA mismatch repair (MMR).^13^ In Enterobacterales, the first steps of MMR are carried out by MutS, MutL, and MutH proteins.^14^ MutS binds to DNA mismatches and recruits MutL. The MutS-MutL-DNA complex binds to and activates latent endonuclease activity of MutH, which is bound to DNA at a neighboring *dam* methylation site. MutH specifically nicks the newly synthesized (unmethylated) DNA strand at the mismatched site, thereby marking the strand containing the erroneously incorporated nucleotide for excisional repair.^13,15,16^ This process is critical for maintaining high fidelity DNA replication. In *Escherichia coli*, loss of function of *mutS, mutL* or *mutH* results in a fixed hypermutation phenotype. Hypermutation is selected for in the harsh environment of infection because it accelerates adaptation by evolution under these conditions. Notably, this includes development of antibiotic resistance via point mutations and by facilitating plasmid acquisition.^17–20^ There are limited data on hypermutation in *K. pneumoniae*, CRKP or other CRE.^21–24^

In this study, we analyzed 11 KPC-producing *K. pneumoniae* isolates collected over approximately 4 years from an immunocompromised patient with chronic GI colonization and recurrent invasive infections. Whole genome sequencing (WGS) revealed emergence of a *mutH* point mutation during CZA treatment, which conferred a V76G substitution in a conserved region of MutH that is important for endonuclease activity. Using clinical and engineered *mutH* mutant strains in *in vitro* and mouse experiments, we demonstrate that *mutH*^V76G^ leads to a hypermutator phenotype, rapid genetic diversification (including development of *bla*_KPC_ mutations that confer CZA resistance and restore meropenem (MEM) susceptibility), increased horizontal plasmid transfer and improved fitness *in vivo*. These findings provide a rare longitudinal view of CRKP evolution in-host and highlight a hypermutator strain as a potent mediator of multidrug resistance and other phenotypes relevant to colonization and disease.

## Results

### *mutH*^V76G^ emerges in longitudinal *Klebsiella pneumoniae* isolates that demonstrate CZA resistance, KPC-3 variant evolution and heightened genomic divergence

We analyzed 11 longitudinal KPC-producing *K. pneumoniae* isolates collected from sites of colonization (C) or invasive disease (D) in an immunocompromised patient. Isolates were categorized into early and late groups [Figure 1, Table 1]. Early isolates (collected within 4 weeks after lung transplantation) included C1 and C2 (i.e., first and second colonization isolates, recovered from serial surveillance rectal swabs), D1 and D2 (i.e., first and second disease isolates, recovered after C1 and C2 and associated with pneumonia and empyema, respectively), and C3 and C4 (recovered from surveillance rectal swabs after D1 and D2). In chronological order, late isolates (collected between 175 weeks (∼3.3 years) and 204 weeks (∼3.9 years) after the initial isolation of C1) were D3 (bacteremia), C5 (rectal surveillance), D4 (tracheal aspirate), C6 (rectal surveillance) and D5 (empyema). WGS confirmed that all isolates belonged to the multilocus sequence type 258 (ST258) lineage and carried *bla*_KPC_ genes.

**Figure 1.**
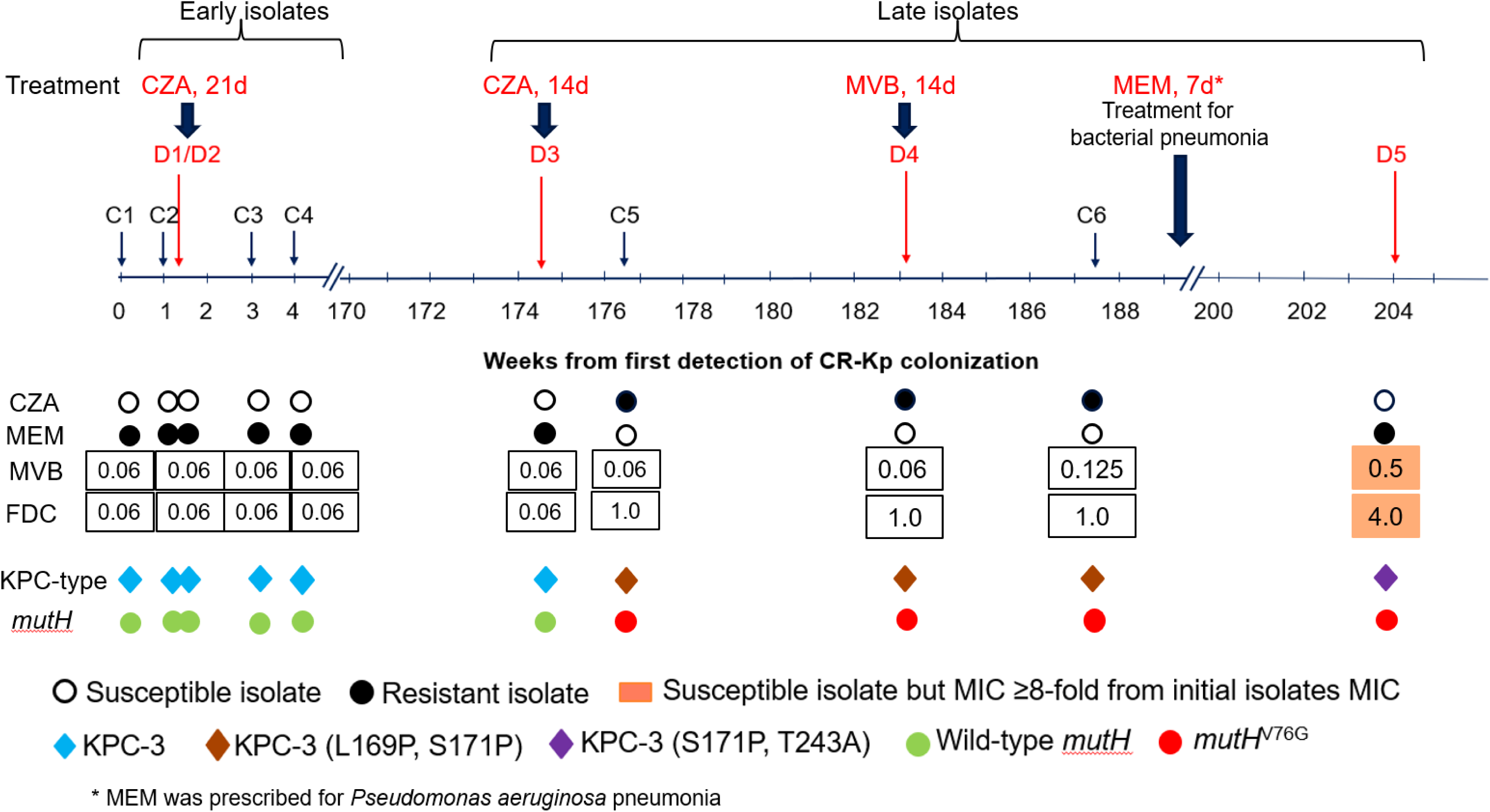
Timeline and features of longitudinal *Klebsiella pneumoniae* isolates. Eleven isolates collected from a lung transplant recipient are included in this study. Six early isolates (C1, C2, D1, D2, C3, C4) were recovered within a 4 week period following lung transplantation. Five late isolates (D3, C5, D4, C6, D5) were recovered > 3 years after C1. Isolates with names prefixed by C (colonization) were recovered from rectal surveillance cultures. Isolates with names prefixed by D were cultured from various sites of suspected disease. Antibiotic treatment regimens are shown above the timeline (start dates noted by black arrows). Antibiotic susceptibility, KPC type and *mutH* genotype are shown below the timeline. MVB and FDC MICs are presented as µg/mL. All isolates were MVB and FDC susceptible, but MICs were 4-64 fold higher against the last isolate (D5) than they were against earlier isolates. CZA: ceftazidime-avibactam; MVB: meropenem-vaborbactam; MEM: meropenem; CFDC: cefiderocol; 21d: 21 days; CR-KP: carbapenem-resistant *K. pneumoniae*; C1: first colonizing isolate; D1: first disease-associated isolate; MIC: minimum inhibitory concentration; KPC: *K. pneumoniae* carbapenemase

**Table 1.**
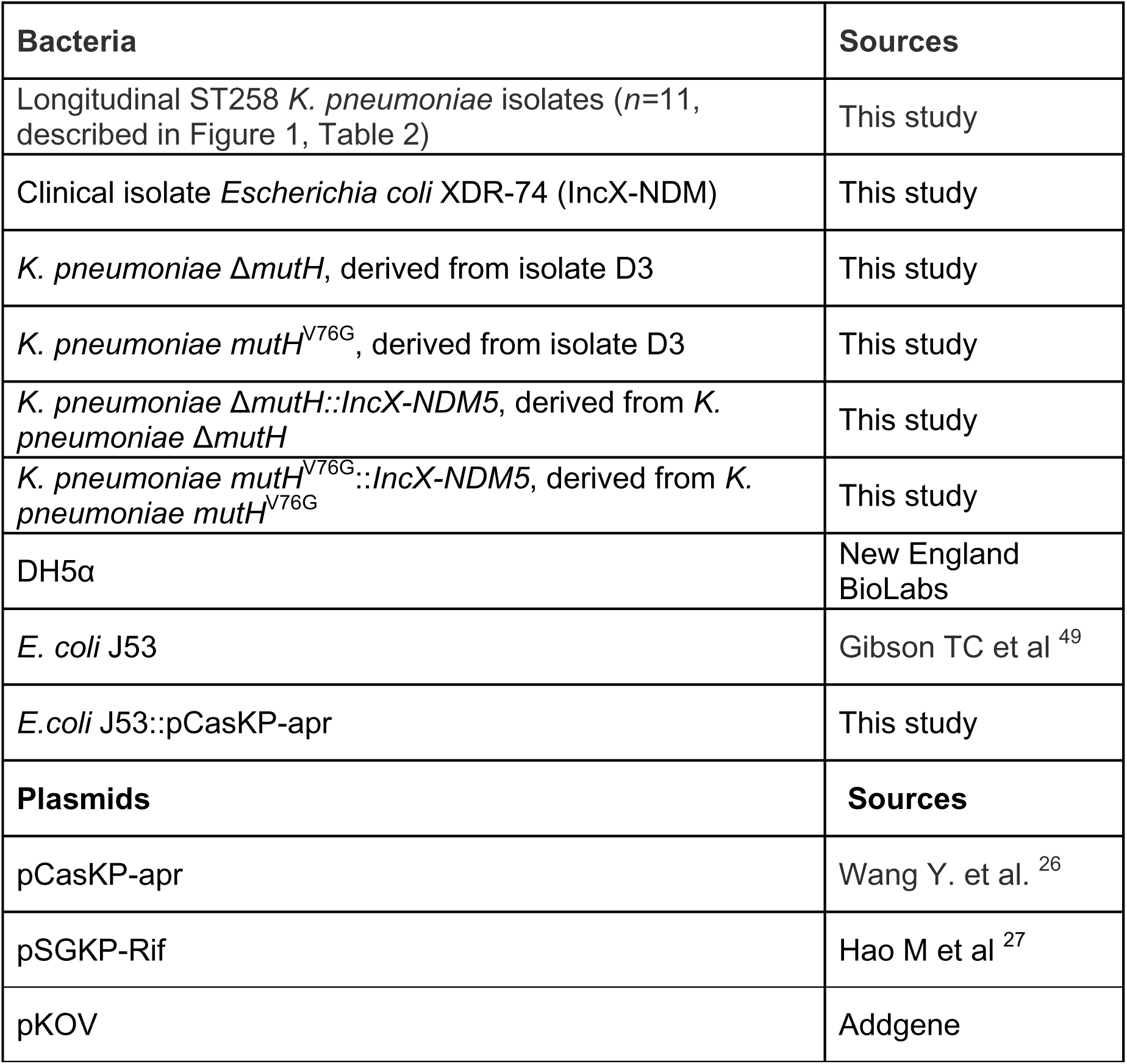
Bacterial strains and plasmids used in this study.

Early isolates and the first late isolate (D3) were MEM resistant (MIC>16 µg/mL) and CZA susceptible (MIC=0.125 µg/mL), and they carried *bla*_KPC-3_. After 2 weeks of CZA treatment for bacteremia caused by isolate D3, late isolates C5, D4 and C6 were CZA resistant but MEM susceptible. Whereas early isolates and D3 harbored wild-type *mutH*, C5, D4 and C6 carried *mutH* with a single nucleotide mutation conferring a MutH^V76G^ substitution [Figure 1; Supplemental Figure S1]. Isolates C5, D4 and C6 acquired *bla*_KPC-3_ mutations conferring L169P and S171P substitutions in the KPC Ω-loop. *bla*_KPC-3_^L169P^ has been implicated in CZA resistance and concurrent restoration of MEM susceptibility.^23^ The final longitudinal isolate (D5) reverted to MEM resistance and CZA susceptibility. D5 maintained *mutH*^V76G^, lost *bla*_KPC-3_^L169P^ and gained *bla*_KPC-3_^T243A^, which confers a substitution outside the Ω-loop that has not been shown to mediate CZA resistance.

All 11 isolates were susceptible to cefiderocol (FDC) and MVB. However, FDC MICs increased 16-fold against late isolates C5, D4 and C6 (MIC=1.0 µg/mL) compared to early isolates and D3 (MIC=0.06 µg/mL). FDC and MVB MICs were 4-fold higher against final isolate D5 (MICs=4.0 and 0.5 µg/mL, respectively) than against penultimate isolate C6 (MICs=1.0 and 0.125 µg/mL, respectively). Late isolates C5 through D5 acquired mutations in *envZ*, which encodes the kinase of a 2-component system involved in iron transport and regulation of outer membrane porins (*envZ*^L19P^ in all isolates; *envZ*^K358I^ also present in D4). In addition, D5 uniquely acquired mutations in the porin *ompK36*^Q301W^ and iron transporter *cirA*^Y294C^. No sequence differences were detected in *PBP3*, regulatory genes (*baeR, ompR*), siderophore uptake genes (*fepA, fhuA, tonB*), or RND efflux pump components (*acrA, acrB, tolC, mexB, oprM*), indicating these loci did not contribute to reduced FDC susceptibility [Table 2].

**Table 2.** Evolution of antibiotic MICs and susceptibility among longitudinal *K. pneumoniae* isolates.

| Isolates | MEM, CZA, MVB susceptibility and MICs | FDC MIC<sup>1</sup> | KPC-3 mutations | OmpK36 mutations | CirA mutations | EnvZ mutations |
| --- | --- | --- | --- | --- | --- | --- |
| C1, C2, D1, D2, C3, C4 (early isolates) | MEM resistant (MIC>16 µg/mL)<br>CZA susceptible (MIC=0.125 µg/mL)<br>MVB susceptible (MIC=0.06 µg/mL) | 0.125 µg/mL | Wild type | Wild type | Wild type | Wild type |
| D3 (late isolate, pre-treatment) | MEM resistant (MIC>16 µg/mL)<br>CZA susceptible (MIC=0.125 µg/mL)<br>MVB susceptible (MIC=0.06 µg/mL) | 0.06 µg/mL | Wild type | Wild type | Wild type | Wild type |
| C5, D4, C6 (late isolates, post-treatment) | MEM susceptible (MIC=0.06 µg/mL)<br>CZA resistant (MIC<br>MVB susceptible (C5, D4 MIC=0.06 µg/mL; C6 MIC=0.125 µg/mL) | 1 µg/mL | L169P, S171P | Wild type | Wild type | L19P, K358I (C5, D4)<br>L19P (C6) |
| D5 (last isolate) | MEM resistant (MIC ≥64 µg/mL)<br>CZA susceptible<br>MVB susceptible (MIC=0.5 µg/mL) | 4 µg/mL | S171P, T243A | Q103W | Y294C | L19P, K358I |

<sup>1</sup>Cefiderocol MICs were increased against C5, D4, C6, D5 but remained within susceptible range. *ompK36*<sup>Q301W</sup>, *cirA*<sup>Y294C</sup>, *envZ*<sup>L19P</sup>, *envZ*<sup>K358I</sup> substitutions are observed uniquely in C5, D4, C6, D5; their functional contributions to increased FDC MICs has not been experimentally confirmed.

Core genome single nucleotide polymorphism (SNP) analysis revealed clear evolutionary structuring of longitudinal isolates. Early isolates were highly clonal, differing by only 2–7 SNPs. The first late isolate (D3) remained closely related to early group (12–15 SNP differences) [Figure 2A]. In contrast, subsequent late isolates (C5, D4, C6, D5) demonstrated more marked divergence, with pairwise distances ranging from 122–264 SNPs compared to early isolates and 69–197 SNPs among themselves [Figure 2B]. Early isolates clustered tightly with D3 by maximum likelihood phylogeny, while later isolates formed progressively more distant branches [Figure 2A]. The findings suggest that acquisition of *mutH*^V76G^ was associated with a transition from long-term genomic stability to heightened within-host diversification.

**Figure 2.**
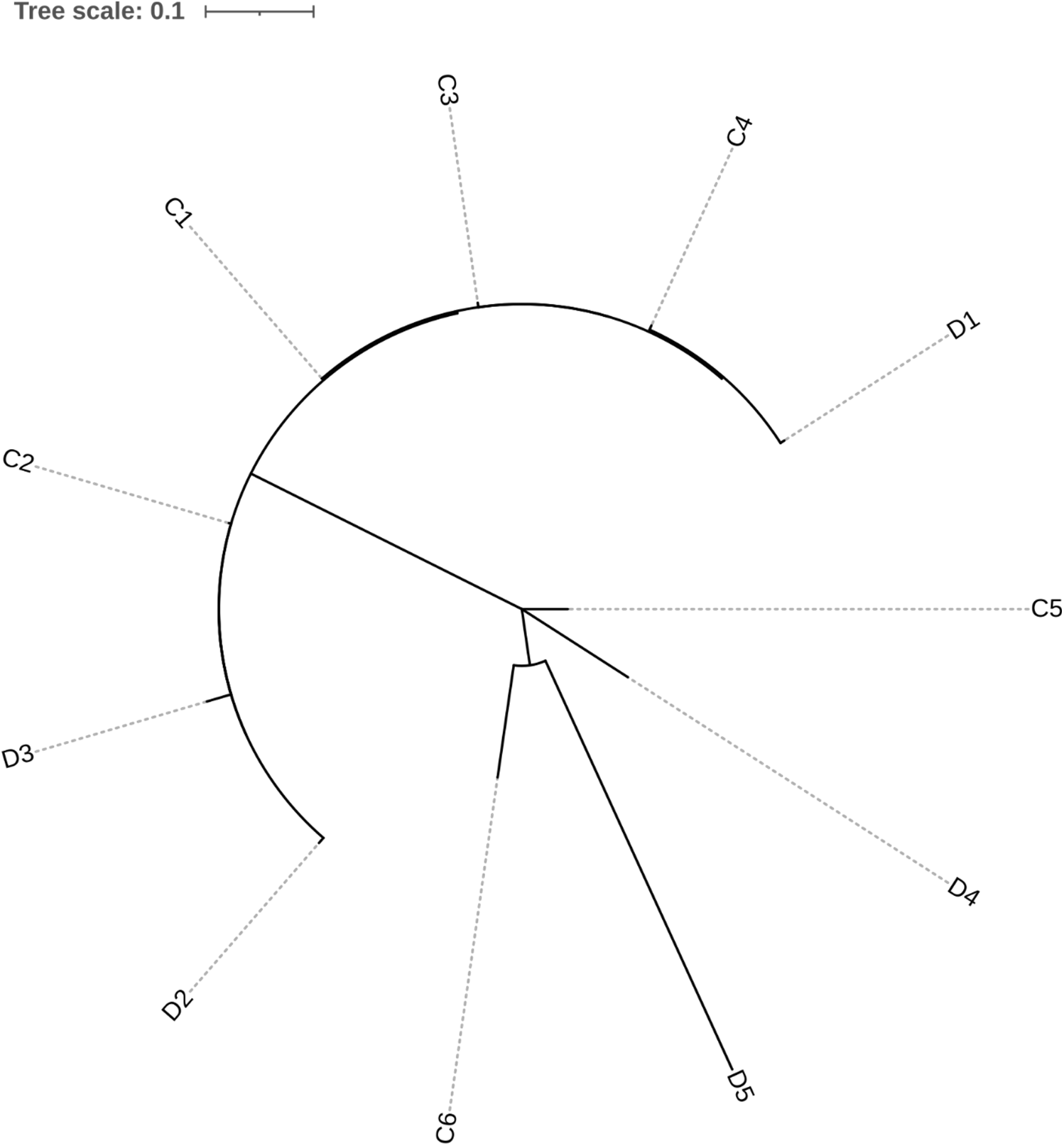
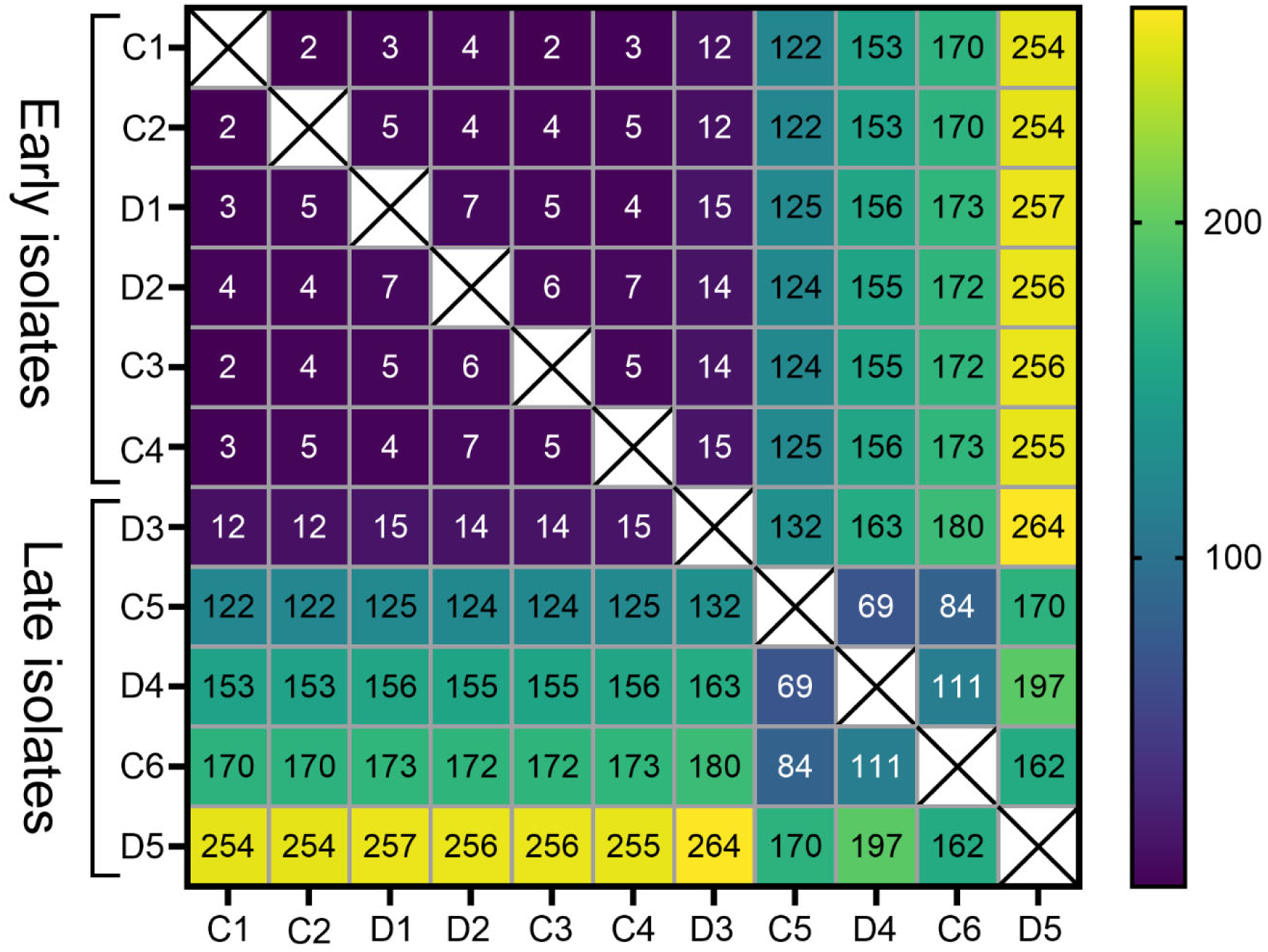
Core genome single nucleotide polymorphism (SNP) phylogeny and SNP distance matrix for longitudinal *K. pneumoniae* isolates. **2A. Core genome SNP-based maximum likelihood phylogenetic tree.** Evolutionary relationships are shown for 11 longitudinal KPC-producing *K. pneumoniae* isolates collected from a lung transplant recipient over almost 4 years. Early isolates (C1, C2, D1, D2, C3, C4) clustered closely with the first late isolate (D3), whereas subsequent late isolates (C5, D4, C6, D5) were progressively more divergent. The tree was built from an alignment of core genome SNPs, with bootstrap support of 100% for every cluster. **2B. Pairwise SNP distance matrix.** The table gives the number of SNP differences between isolates. The color gradient reflects genomic divergence, with dark purple indicating closely related isolates (low SNP distances) and yellow indicating greater divergence (high SNP distances). Early isolates (C1, C2, D1, D2, C3, C4) and the first late isolate (D3) were highly similar, differing by only 2–15 SNPs. In contrast, later isolates carrying *mutH^V76G^* (C5, D4, C6, D5) showed progressively greater divergence from the earlier cluster (122–264 SNPs) and from one another (69–197 SNPs), consistent with a hypermutator phenotype driving accelerated within-host evolution.

### *mutH*^V76G^ confers a conserved motif substitution that results in hypermutable phenotypes and increased horizontal transfer of plasmids

*K. pneumoniae* MutH shares 82.5% sequence identify with *E. coli* MutH, for which a crystal structure has been solved.^25^ Alignment of *K. pneumoniae* MutH to other MutH family members shows that V76 is part of a highly conserved five residue sequence motif (GVELK) that includes two active site residues (E77, K79), two residues that contribute to the hydrophobic core of the N-terminal subdomain (V76, L78), and a glycine (G75) predicted to be important for structural stability [Figure 3].^25^

**Figure 3.**
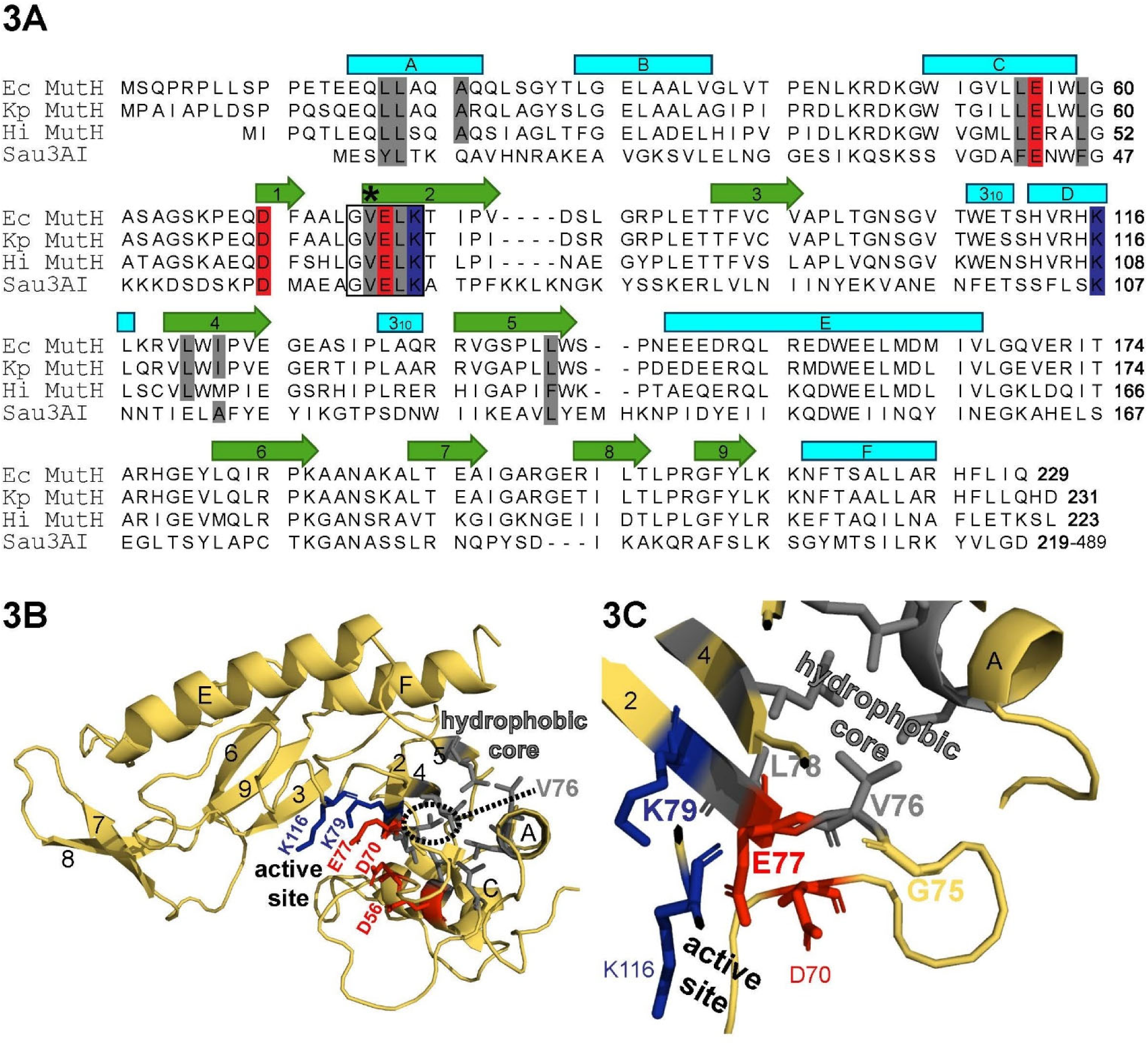
Structural analysis of V76G substitution on MutH. **3A.** Sequence alignment of the MutH family), including MutH from *Escherichia coli* (Ec), *Klebsiella pneumoniae* (Kp), *Haemophilus influenzae* (Hi), and Sau3AI from *Staphylococcus aureus*. Alignment is adapted from Ban and Yang, *EMBO J.* 17(5):1526–1534 (1998). The conserved GVELK motif is boxed and the position of residue V76 is indicated with an asterisk. Secondary-structure elements from the *E. coli* MutH crystal structure are shown above the alignment, where cyan boxes denote α-helices and green arrows denote β-strands. **3B.** Ribbon diagram of the published MutH structure from *E. coli* (pdb: 2AZO; Ban and Yang, EMBO J. 17(5):1526-1534, 1998) and **3C** the region surrounding V76. Residues of interest are drawn as sticks. Active site residues are shaded red (acidic) or blue (basic). Residues comprising a hydrophobic core between α-helices A and C, and β-sheets 2, 4, and 5 are shaded grey. Residues 75-79 of the conserved GVELK motif (β-sheet 2) are bolded in panel C. The analysis shows that the nonpolar sidechain of V76 points into a hydrophobic core, which likely assists in stabilizing the relative positioning of the neighboring active site residues to allow for catalysis. V76G substitution is expected to disrupt catalysis due to loss of this stabilizing hydrophobic interaction.

To assess the functional impact of this substitution, we examined the hypermutator phenotype associated with *mutH*^V76G^. Late isolates carrying *mutH*^V76G^ exhibited significantly elevated rifampin mutational frequencies (RFP) *in vitro* compared to wild type *mutH* isolates (log_10_ mean ± standard error of mean (SEM): -5.75±0.09 *vs.* - 7.48±0.08, *p* < 0.00001) [Figure 4A,]. RFP >log_10_ -6 is typically taken as evidence of a hypermutable isolate.^23^ In 14-day serial passage experiments with escalating MEM concentrations and fixed vaborbactam (8 µg/mL) concentration *in vitro*, representative *mutH*^V76G^ isolates (C5, D5) developed MVB resistance by passage 3 and attained MICs >256 µg/mL by passage 8. In contrast, representative *mutH* wild type strains (C3, D3) remained susceptible after 14 passages [Figure 5A, upper panel].

**Figure 4.**
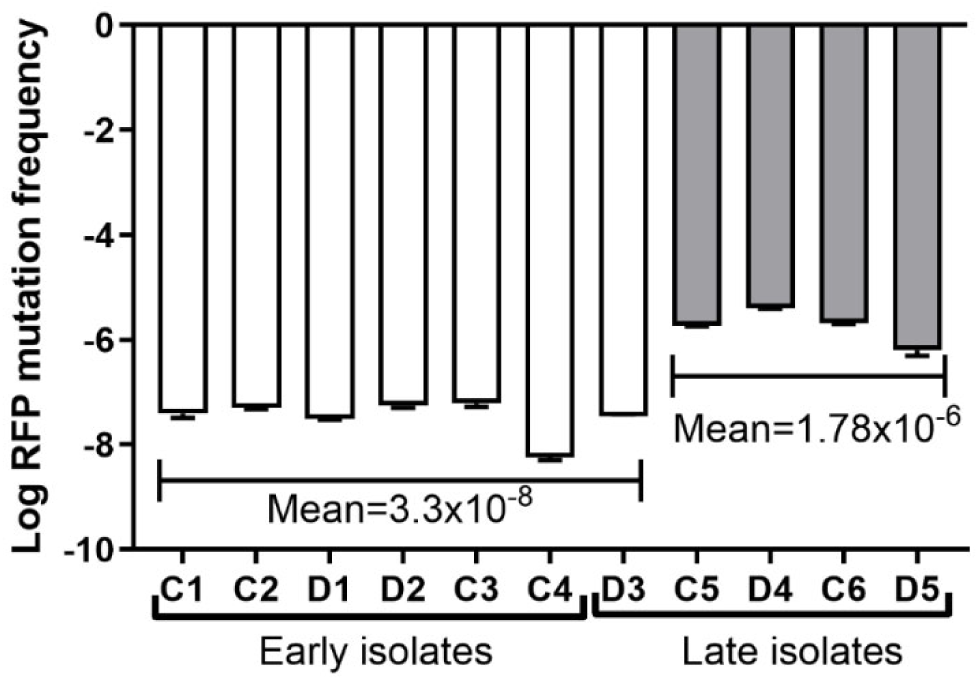
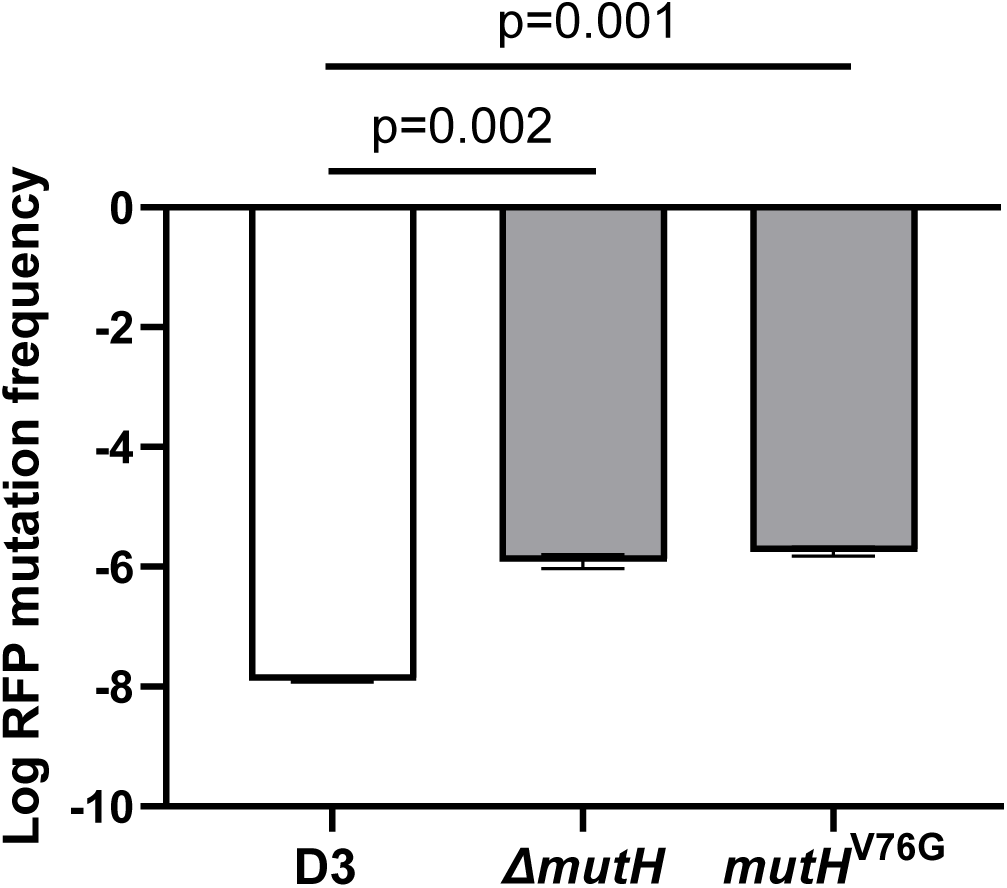
Evaluation of the contribution of *mutH* to rifampin mutational frequency (RFP). **4A** shows RFPs of clinical isolates, and **4B** shows results from engineered isogenic strains differing only in the *mutH* allele (wild type, Δ*mutH*, and *mutH*^V76G^). Engineered mutants were generated from *mutH* wild-type isolate D3 using a *Klebsiella*-specific CRISPR–Cas9 platform. Data represent three independent experiments. **4A.** Log₁₀ RFP of 11 longitudinal clinical isolates. *mutH* mutant isolates exhibited significantly higher log₁₀ RFP than wild-type isolates (*p* < 0.0001). White and grey bars denote *mutH* wild-type and mutant isolates, respectively. **4B.** Log_10_ RFP of engineered isogenic strains. Both engineered mutants demonstrated higher RFP than the D3 parent strain (Δ*mutH*, –5.91 ± 0.12; *mutH*^V76G^, –5.74 ± 0.08; vs. D3, –7.89 ± 0.03; *p*= 0.002 and 0.001, respectively; two-way ANOVA with Sidak’s multiple comparisons test).

**Figure 5.**
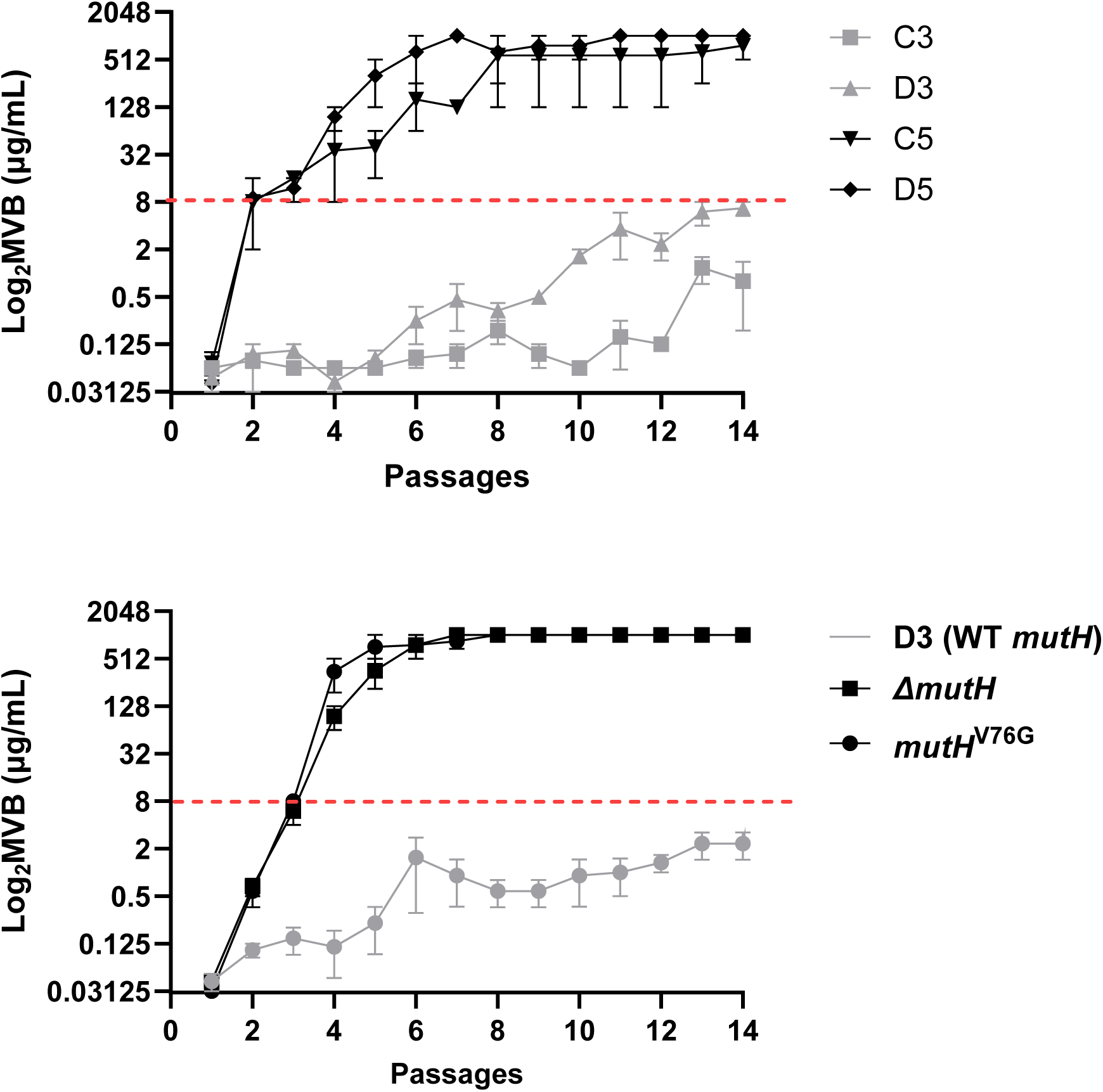
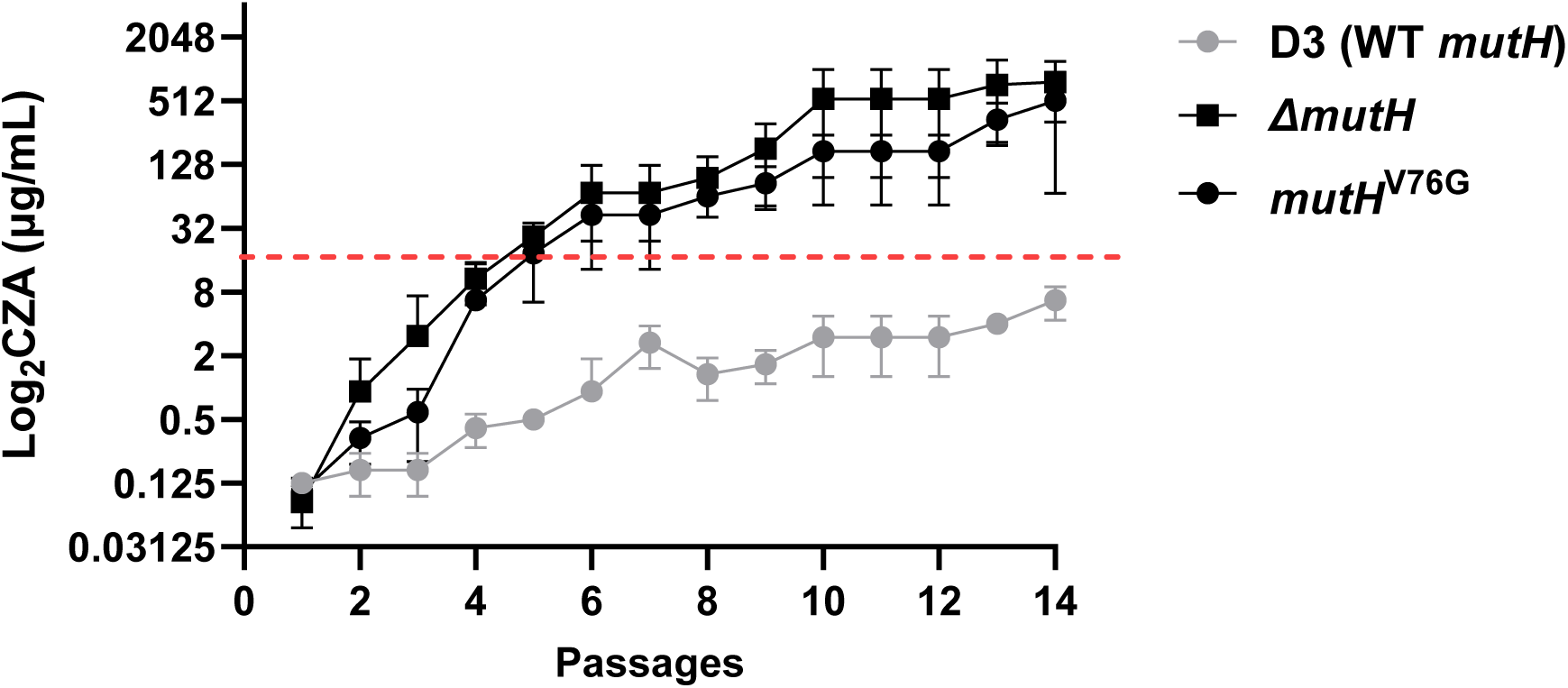
Impact of *mutH* on the evolution of antibiotic resistance during serial passage in meropenem-vaborbactam (MVB) and ceftazidime-avibactam (CZA) *in vitro*. Fourteen serial passages were performed using escalating concentrations of MVB (Fig. 5A) or CZA (Fig. 5B). For MVB, meropenem (0.03–1,024 µg/mL) was combined with a fixed vaborbactam concentration of 8 µg/mL; for CZA, ceftazidime (0.03–1,024 µg/mL) was combined with fixed avibactam at 4 µg/mL. CLSI nonsusceptibility breakpoints for MVB (≥8/8 µg/mL) and CZA (≥16/4 µg/mL) are indicated by red dashed lines. Data are from three independent experiments. **5A (upper).** Serial passage of four clinical isolates in MVB, two of which harbored wild-type *mutH* (C3, D3) and two of which harbored *mutH*^V76G^ (C5, D5). Mutant isolates developed MVB resistance by passage 3, with MICs exceeding 256 µg/mL by passage 8, whereas wild-type isolates (C3, D3) remained susceptible through all 14 passages. **5A (lower):** Serial passage of the *mutH* wild-type isolate D3 and two engineered mutants (Δ*mutH* and *mutH*^V76G^) in MVB. Engineered *mutH* mutants exhibited more rapid increases in MVB MICs than the D3 parent strain, achieving resistance by passage 3. **5B.** Serial passage of engineered *mutH* mutants in CZA. *mutH* mutants developed elevated CZA MICs more rapidly than wild-type D3 and became resistant after 5 passages. CZA passage experiments were not performed for clinical isolates because *mutH*^V76G^ isolates were already CZA-resistant.

We constructed isogenic Δ*mutH* (*mutH* gene disruption) and *mutH*^V76G^ strains from isolate D3 using a *Klebsiella*-specific CRISPR-Cas9 genome editing platform.^26^ WGS confirmed no off-target effects.^27^ Mutants had 106-fold (Δ*mutH*) and 157-fold (*mutH*^V76G^) higher RFP than D3 (mean log_10_: -5.91±0.12 and -5.74±0.08, respectively, *vs.* -7.94 ± 0.02) [Figure 4B]. In MVB passage experiments, mutant strains became MVB resistant after passage 3, while D3 remained susceptible through passage 14 [Figure 5A, lower panel]. In experiments with escalating ceftazidime concentrations and fixed avibactam concentration (4 µg/mL), mutant strains developed CZA resistance by passage 5, while D3 remained susceptible through passage 14 [Figure 5B].

To study conjugative transfer, we first used *E. coli* XDR-74 with IncX3 plasmid carrying *bla*_NDM-5_ [Supplemental Figure S2] as the donor strain and engineered Δ*mutH* and *mutH*^V76G^ mutants and D3 as recipients.^28^ Acquisition of IncX3 with *bla*_NDM-5_ by Δ*mutH* and *mutH*^V76G^ mutants was significantly greater than by D3 (conjugation frequencies ∼1,820-fold (3.36-log) and 930-fold (2.97-log) higher, respectively; *p* < 0.0001) [Figure 6A]. Next, we used *E. coli* J53 harboring pCasApr as recipient and Δ*mutH*, *mutH*^V76G^ or D3 carrying IncX3 with *bla*_NDM-5_ as donor. Δ*mutH* and *mutH*^V76G^ strains exhibited modest but significant increases in conjugation frequency compared to D3 (∼2-fold and ∼2.75-fold, respectively; *p*=0.04 and 0.01) [Figure 6B]. Therefore, *mutH* mutant strains significantly increased donor and, in particular, recipient efficiency during conjugative plasmid transfer.

**Figure 6.**
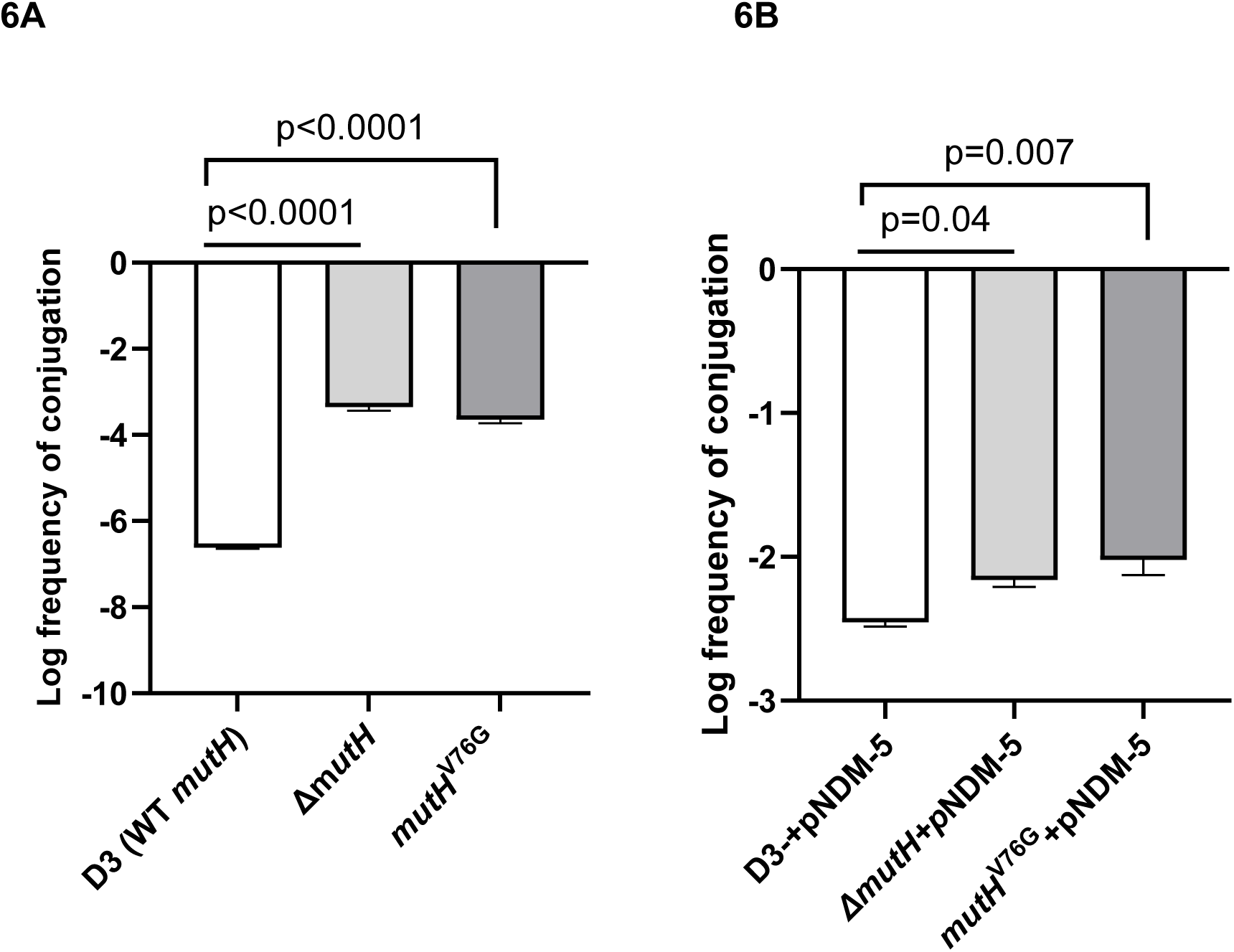
*mutH* impact on conjugation frequency. Conjugation frequencies were calculated based on recovery of recipient strain transconjugants, reflecting efficiency of plasmid transfer from donor strains. **6A. Conjugation frequency for recipient *K. pneumoniae* strains harboring wild type or engineered mutant *mutH*.** The donor was *E. coli* clinical isolate XDR-74 carrying the IncX3 plasmid with *bla*_NDM-5_. Conjugation frequencies of Δ*mutH* (log_10_ –3.36±0.08) and *mutH*^V76G^ (log_10_ –3.65±0.08) isogenic strains were significantly higher than that of wild type *mutH* parent isolate D3 (log_10_ –6.62±0.02; *p* <0.0001 for both comparisons, one-way ANOVA with Dunnett’s multiple comparisons test). **6B. Conjugation frequency for donor *K. pneumoniae* strains harboring wild type (isolate D3) or mutant *mutH* (Δ*mutH*, *mutH^V76G^*).** Donor strains carried the IncX3 plasmid with *bla*_NDM-5_. The recipient was *E. coli* J53 carrying pCasApr. Δ*mutH* (log_10_ –2.16±0.05) and *mutH^V76G^* (log_10_ –2.02±0.10) strains exhibited significantly higher conjugation frequency than did wild type *mutH* parent strain D3 as donor (log_10_ –2.46±0.28; *p*=0.04 and *p*=0.007, respectively; one-way ANOVA with Dunnett’s multiple comparisons test).

### *mutH*^V76G^ enhances fitness and promotes emergence of CZA and MVB resistance in vivo

We immunocompromised CD1 mice (6 per group) with intraperitoneal cyclophosphamide (150 mg/kg 4 days prior and 100 mg/kg 1 day prior to infection) and subcutaneous cortisone (20 mg/kg 4 days prior and 1 day prior). We then intravenously infected mice with engineered isogenic *mutH* mutant strains or D3 (5×10⁴ CFU/mouse) and quantified bacterial burdens within livers, kidneys and spleens on day 3. In each organ, burdens of Δ*mutH* were significantly higher than those of D3 (mean±SEM log_10_ CFU/g, liver: 5.70±0.84 *vs.* 2.05±0.48, *p*=0.014; kidney: 4.30±0.51 *vs.* 1.66±0.34, *p*=0.014; spleen: 5.84±0.53 *vs.* 2.22±0.72, *p*=0.02) [Figure 7A]. Burdens of the *mutH*^V76G^ strain were significantly higher than those of D3 in liver (7.40±0.10 *vs.* 2.05±0.48, *p*=0.0004) and spleen (5.64 ± 0.28 *vs.* 2.22±0.72, *p*=0.008). Burdens of Δ*mutH* and *mutH*^V76G^ did not differ significantly in any organ.

**Figure 7.**
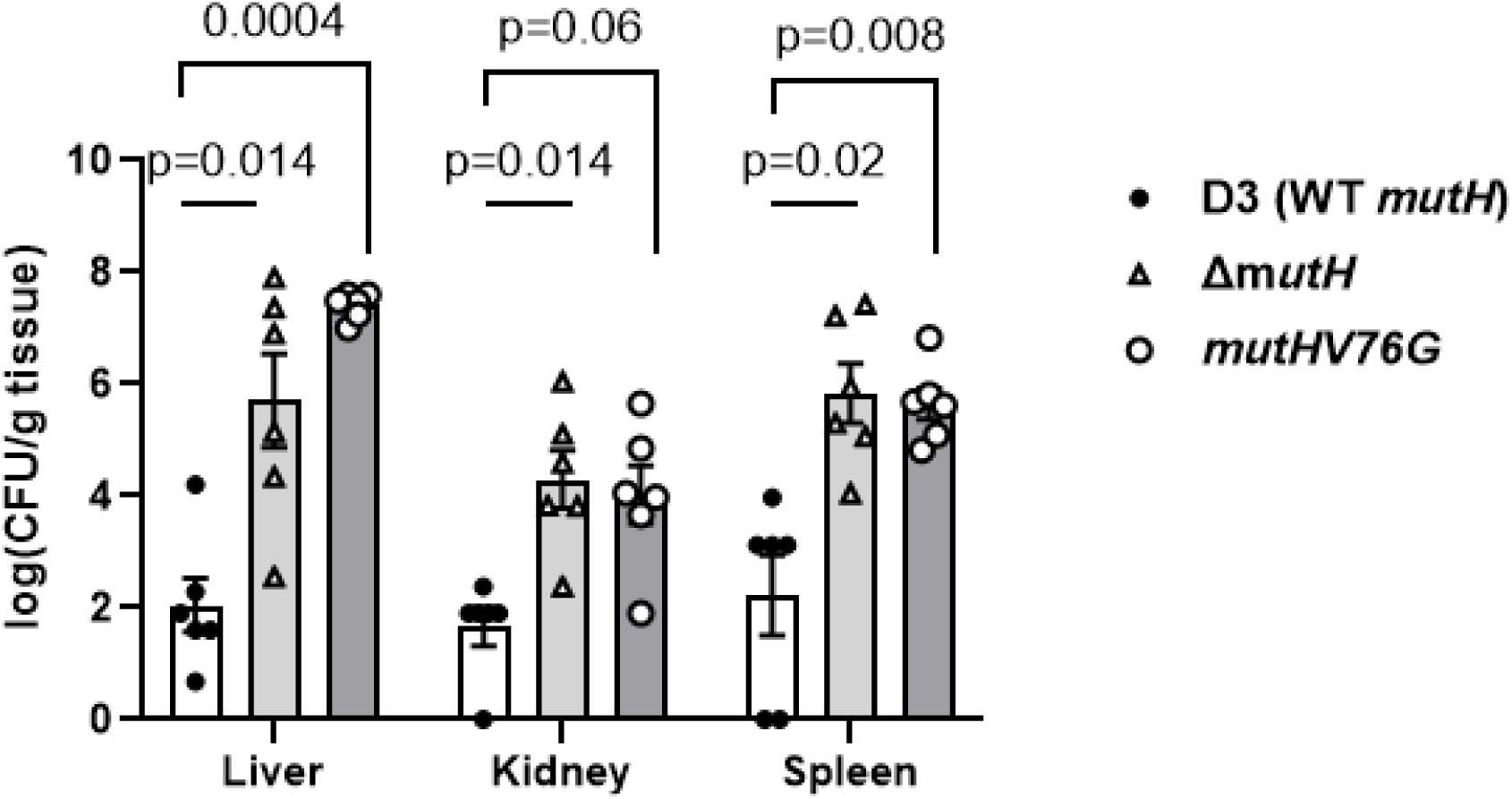
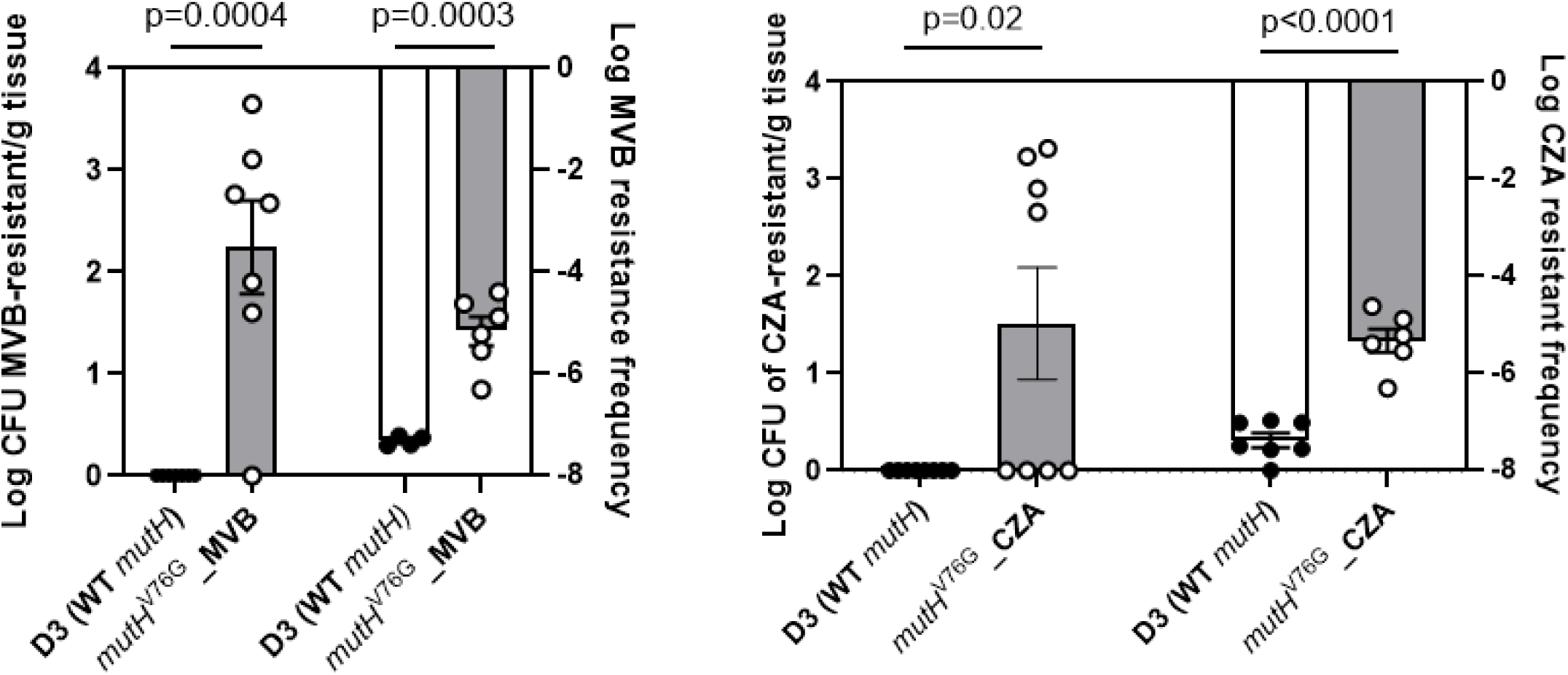
*mutH*^V76G^ impact on *K. pneumoniae* fitness *in vivo* and emergence of resistance during MVB and CZA treatment. **7A. *mutH* impact on strain fitness.** Mice were infected intravenously with 5×10⁴ CFU/mouse of wild type *mutH* isolate D3 or engineered *mutH* mutant strains (Δ*mutH* or *mutH*^V76G^). Mice were sacrificed on day 3 and bacterial burdens were quantified in livers, kidneys and spleens. In livers, mean±SEM log_10_ CFU/g was 2.05±0.48 (D3), 5.70±0.84 (Δ*mutH*, *p*=0.014) and 7.40 ± 0.210 (*mutH*^V76G^, *p*=0.0004). In kidneys, mean burdens were 1.66±0.34 (D3), 4.30±0.54 (Δ*mutH*, *p*=0.014) and 4.02±0.51 (*mutH*^V76G^, *p*=0.06) log_10_ CFU/g. In spleens, mean burdens were 2.22±0.72 (D3), 5.84±0.53 (Δ*mutH*, *p*=0.02) and 5.64 ± 0.28 (*mutH^V76G^*, *p*=0.008) log_10_ CFU/g. Statistical analysis was performed using two-way ANOVA with Turkey’s multiple comparisons test. **7B. *mutH* impact on emergence of MVB or CZA resistance.** Mice were infected intravenously with 1×10^5^ CFU/mouse of isolate D3 or the isogenic *mutH^V76G^* strain. The *mutH^V76G^* strain was chosen since this mutation rather than Δ*mutH* arose in clinical isolates. MVB and CZA MICs against D3 and the *mutH^V76G^* strain were 0.125 µg/mL and 0.25 µg/mL, respectively. Mice were treated with MVB (50 mg/kg, intraperitoneal, every 8 hours) or CZA (37.5 mg/kg intraperitoneal every 8 hours) starting 4 hours post-infection. On day 3, mice were sacrificed, and bacterial burdens in liver, kidney, and spleen were quantified. Homogenates were plated on LB agar with and without 4xMIC of the respective antibiotic. Log_10_ resistance frequency indicates the proportion of recovered bacteria within a population and is calculated as: log_10_ (bacterial count on 4xMIC plates/bacterial count on no antibiotic plates). Left panel: Log_10_ CFU/g of strains in tissues from MVB treated mice, as identified by growth on plates containing 4xMIC MVB. Strains were recovered exclusively from the livers of mice infected with the *mutH^V76G^* mutant (2.24 ± 0.46 log_10_ CFU/g enumerated on MVB-containing plates). No strains were isolated from MVB treated D3-infected mice. Log_10_ resistance frequencies for D3 and *mutH^V76G^* were – 7.31±0.05 and –5.1 ±0.28, respectively (*p*= 0.0003). All isolates recovered from MVB-containing plates were confirmed to have MVB MICs ≥ 16 µg/mL. Right panel: Log_10_ CFU/g of strains in tissues from CZA treated mice, as identified by growth on plates containing 4xMIC CZA. Strains were recovered exclusively from the livers of mice infected with the *mutH^V76G^* mutant (1.51±0.24 log_10_ CFU/g enumerated on CZA-containing plates)). No strains were isolated from CZA treated D3-infected mice. Log_10_ recovery frequencies for D3 and *mutH^V76G^* were – 7.37± 0.15 and –5.33±0.24, respectively (*p*<0.0001). Among 16 randomly selected isolates from CZA-containing plates, 7 were confirmed to have CZA MICs ≥ 16 µg/mL.

In other experiments, we intravenously infected mice treated mice with the *mutH*^V76G^ strain or D3 (1×10^5^ CFU/mouse) and treated them beginning 4 hours later with MVB or CZA [Figure 7B]. For D3-infected mice, no colonies were recovered from organ homogenates cultured on plates containing the respective antibiotic at 4xMIC. In contrast, homogenized livers of *mutH*^V76G^-infected mice cultured on such plates after MVB or CZA treatment yielded 2.24±0.46 and 1.51±0.24 mean log_10_ CFU/g (±SEM), respectively. Corresponding MVB recovery frequencies of *mutH*^V76G^ and D3 were log_10_ –5.16±0.28 and –7.31±0.05 (*p*=0.0003) [Figure 7B, left panel]. CZA recovery frequencies were log_10_ –5.33±0.24 and -7.37±0.15, respectively (*p*<0.0001). [Figure 7B, right panel].

All 57 isolates recovered from MVB-containing plates were resistant to MVB *in vitro* (MICs ≥16 µg/mL). Sequencing of *ompK36* from 5 randomly selected MVB-resistant isolates identified mutations predicted to impair OmpK36 function, including nonsense substitutions (Q92STOP (1 strain), Q76STOP (2 strains)) and missense variants (L361P, L363P (1 strain each)). The 5 isolates also showed significantly reduced *ompK36* transcript levels compared to D3, as determined by qRT-PCR, consistent with reduced OmpK36 protein levels on outer membrane analysis [Supplementary Figure S3].

Among 16 isolates selected randomly from CZA-containing plates, 7 were resistant to CZA (MICs ≥16 µg/mL). Sequencing identified *bla*_KPC-3_^A172T^ (*n*=4) and *bla*_KPC-3_^R164H^ (*n*=2), both of which have been linked to CZA resistance.^29,30^

## Discussion

We demonstrate that KPC-producing *K. pneumoniae* isolates causing long-term colonization and recurrent infections of a lung transplant recipient developed a hypermutator phenotype due to emergence of a V76G substitution in MMR endonuclease MutH. Following CZA treatment, emergent *mutH*^V76G^-carrying isolates manifested CZA resistance and restored MEM susceptibility, accompanied by rapid accumulation of core genome SNPs and mutations in the KPC Ω-loop, OmpK36 porin and other proteins that have been previously linked to changes in antibiotic susceptibility. Using isogenic mutant *K. pneumoniae* strains derived from a wild-type *mutH* clinical isolate, we confirmed that *mutH*^V76G^ accelerates acquisition of antibiotic resistance *in vitro* and *in vivo*, promotes transfer and acceptance of antibiotic resistance-conferring plasmids, and increases strain fitness during mouse infections. Our findings suggest that hypermutability stemming from impaired MMR provides *K. pneumoniae* with substantial adaptive advantages in the face of antibiotic exposures and during prolonged within-host residence.

Functional defects in Mut proteins lead to accumulation of genome mutations that can impact antibiotic resistance, stress responses and other adaptive traits.^31^ To date, mutations in *mutH*, *mutS* and *mutL* that confer hypermutator phenotypes have been described rarely among ST258 CRKP or other clinical *K. pneumoniae* isolates.^21,22,24,32^ In a search of National Center for Biotechnology Information WGS databases, 3.1% of *K. pneumoniae* carried a missense mutation in *mutS*.^22^ The published data likely underestimate the prevalence of MMR-deficient, hypermutator *K. pneumoniae* because the phenotype is uncommonly tested, single rather than multiple isolates are typically characterized from a given culture or patient, most studies focus on Mut proteins (and MutS in particular) and do not consider downstream MMR components, and research is biased toward isolates from acute, sterile site infections instead of colonization or chronic infections in which mutational frequency is higher.^10,21,31,33^

*mutH*^V76G^ variants have not been described previously. However, comparison to other MutH family members’ sequences and available structures suggests the mechanism by which V76G substitution causes loss of function [Figure 3A].^34^ MutH is composed of two subdomains, referred to as the N-arm and C-arm; the former contains the active site residues. The nonpolar sidechain of V76 points into the hydrophobic core of the N-arm [Figure 3B and 3C], which likely assists in stabilizing the relative position of the neighboring active site residues to allow for metal ion coordination and/or catalysis.^34^ V76G substitution is expected to disrupt catalysis due to loss of this stabilizing hydrophobic interaction.

The first 7 longitudinal isolates in this study were KPC-producing *K. pneumoniae* with wild type *mutH* that were collected over ∼3.3 years. These isolates differed by 2-15 SNPs, consistent with reported *K. pneumoniae* evolution rates of 3.8–10 SNPs/genome/year.^6,35,36^ Following a 2-week CZA treatment course for bacteremia by the seventh isolate (D3), *mutH*^V76G^ hypermutator isolate C5 was recovered from a rectal surveillance culture. C5 differed from previous isolates by 122-132 SNPs, far exceeding the expected within-host evolutionary rate, and showed a mutational spectrum consistent with MMR deficiency. The 3 subsequent isolates collected over the next 21 weeks were also *mutH*^V76G^ mutants and exhibited rapidly increasing divergence, differing from C5 and earlier isolates by 69-170 and 153-264 SNPs, respectively. Despite their divergence, these late isolates retained the same ST258 background, *bla*_KPC-3_ backbone, and conserved chromosomal and plasmid features. The sudden increase in genome-wide SNPs coinciding with *mutH* inactivation and large pairwise distances among late isolates strongly suggest a single hypermutator lineage.^21,32^ Within-host evolution of a single strain driven by *mutH*^V76G^-mediated hypermutation is further supported by the temporal succession of isolates and correlations between antibiotic exposure, MICs and acquisition of resistance-conferring mutations. Nevertheless, we acknowledge that we cannot definitively exclude that the patient acquired a new, highly related ST258 strain, although this scenario is less parsimonious.

Serial changes in antibiotic MICs against *mutH*^V76G^-carrying clinical isolates were consistent with specific emergent mutations. CZA resistance first developed in isolate C5, accompanied by reversion from carbapenem resistance to susceptibility. The *bla*_KPC-3_^L169P^ substitution, acquired by C5 and maintained in isolates D4 and C6, mediates these phenotypes by disrupting avibactam binding, enhancing ceftazidime hydrolysis and attenuating carbapenemase activity.^36,37^ The final isolate (D5), recovered following MEM treatment for bacterial pneumonia, was again CZA susceptible and carbapenem resistant, likely due to loss of *bla*_KPC-3_^L169P^. D5 also acquired a *bla*_KPC-3_^T243A^ substitution outside the Ω-loop. Mutations of T243 have been associated with increased carbapenem MIC, but they have inconsistent impact on CZA susceptibility.^38,39^ D5 additionally harbored an *ompK36*^Q301W^ substitution. Although OmpK36 alterations frequently contribute to carbapenem resistance and have been reported in CRKP strains with elevated MVB MICs, the functional significance of the Q301W mutation is unknown. Susceptibility to FDC, a siderophore cephalosporin structurally related to ceftazidime, was also diminished among CZA-resistant isolates. Cross-resistance to CZA and FDC in clinical KPC-producing *K. pneumoniae* has been attributed to combinations of KPC variants, OmpK36 alterations, and mutations in EnvZ and/or CirA, which may impact β-lactamase remodeling, porin diameter, siderophore transport and drug influx/efflux.^40–42^ However, the individual contributions of the specific mutations identified here have not been confirmed experimentally.^41,43,44^

Observations from our clinical isolates were validated with an engineered isogenic *mutH*^V76G^ strain, which displayed markedly accelerated resistance evolution and enhanced adaptability under antibiotic pressure. The mutant strain exhibited increased RFP compared to isogenic parent isolate (D3) and rapidly acquired resistance to MVB and CZA during serial *in vitro* passage and during invasive mouse infections. MVB-treated mice infected with the *mutH*^V76G^ strain accumulated *K. pneumoniae ompK36* mutations (Q76STOP, L361P, L363P) that truncate the porin or restrict its channel size, while CZA-treated mice developed KPC Ω-loop substitutions (A172T, R164H) that influence β-lactamase substrate binding and mediate CZA resistance.^29–31,45^ These findings recapitulated the temporal evolution of MICs and mutations in clinical isolates and confirmed that *mutH*^V76G^-driven hypermutation promotes subsequent acquisition of mutations that attenuate antibiotic susceptibility. In absence of antibiotic treatment, the *mutH*^V76G^ strain was more fit than D3 within mouse livers and kidneys following hematogenous dissemination. The data align with prior findings that hypermutator *E. coli*, *P. aeruginosa* and *S. enterica* were better able than non-hypermutators to survive immune and antibiotic stresses *in vivo.*^9,13,46–49^

*mutH*^V76G^ hypermutation also amplified bidirectional horizontal gene transfer. Conjugation assays revealed that the isogenic *mutH*^V76G^ mutant had 900-fold (∼3-log) increase in acquisition of an IncX3 plasmid carrying *bla*_NDM-5_ from an *E. coli* strain, consistent with the role of MMR in rejecting divergent DNA. Plasmid donation by the *mutH*^V76G^ mutant increased modestly (∼2-fold), in keeping with a more limited and asymmetric influence of MMR in regulating horizontal gene transfer. Overall, our data suggest that *K. pneumoniae* hypermutators may promote resistance directly by facilitating adaptive mutations under antibiotic selection, and indirectly by potentiating acquisition and, to a lesser extent, transfer of resistance gene-bearing plasmids in the bacterial population.^9,13,50^ In assays of conjugation and other phenotypes, results with the *mutH*^V76G^ mutant were similar to those with an isogenic *ΔmutH* strain. Therefore, the V76G substitution appears to largely nullify MutH function.

The strength of this study is its high-resolution view of long-term within-host bacterial evolution. Our data capture emergence of hypermutation, antibiotic resistance and enhanced adaptability in *K. pneumoniae* under clinical selective pressures. Limitations include the single-patient scope, which may not reflect the full diversity of hypermutator phenotypes, and the fact that while mutations in *cirA, envZ,* and *ompK36* coincided with higher FDC or carbapenem MICs, their functional contributions are unproven. We cannot determine whether *mutH*^V76G^ or KPC Ω-loop mutations occurred first, as they were first detected together in isolate C5. However, the sharp increase in SNP accumulation beginning with C5 suggests that hypermutation likely preceded or coincided with acquisition of CZA-resistance mutations, providing the mutational substrate for rapid selection. The data strongly support a model in which *mutH*^V76G^-driven hypermutation accelerated evolution of CZA resistance rather than arising as a consequence of it.

In conclusion, this study demonstrates that *mutH*^V76G^ drives hypermutation, accelerates resistance development, and amplifies horizontal plasmid and resistance gene transfer in CRKP. The ability of *mutH* mutants to rapidly evolve intrinsic resistance and acquire extrinsic resistance determinants underscores their clinical threat. A crucial goal for future studies will be to systematically define the prevalence and clinical impact of *K. pneumoniae* hypermutator strains during infections. Ideally, such studies will screen multiple isolates from individual cultures and patients for hypermutator phenotypes, identify mutations in *mut* and other genes involved in MMR, and correlate phenotypes and genotypes with responses to antibiotic treatment and patients’ outcomes.

## Materials and methods

### Clinical isolates, strains and antimicrobial susceptibility testing

Clinical isolates, bacterial strains and plasmids used in this study are listed in Table 1. Unless otherwise specified, bacteria were cultured in 4 mL of Mueller–Hinton (MH) broth at 37°C with shaking at 180 rpm. Eleven *Klebsiella pneumoniae* isolates were collected from an immunocompromised patient enrolled in the CREST study (CRE and MDR-E carriage among solid organ transplant recipients),^6^ which was approved by the University of Pittsburgh Institutional Review Board. Both colonizing- and disease-causing isolates were preserved at –80°C. *E. coli* XDR-74 carrying an IncX3 plasmid with *bla*_NDM-5_ was obtained from the UPMC XDR Pathogen laboratory. Species identification was performed by MALDI-TOF MS (Bruker). Antimicrobial susceptibility testing was conducted by broth microdilution according to Clinical and Laboratory Standards Institute (CLSI) guideline,^51^ and MICs were interpreted using CLSI or FDA breakpoints.

### Whole-genome sequencing and phylogenetic analysis

Genomic DNA was extracted from overnight cultures using the Wizard Genomic DNA Purification Kit (Promega, Madison, WI). Sequencing libraries were prepared with the Nextera DNA Library Prep Kit and sequenced on an Illumina HiSeq platform (150 bp paired-end reads). Reads were quality-trimmed with Trimmomatic v0.39 and assembled *de novo* using SPAdes v3.15. Genome annotation was performed with Prokka. Multi-locus sequence typing and resistance gene detection were conducted with AMRFinderPlus v3.9 and Kleborate v2.0.1. Comparative genomics of ST258 *K. pneumoniae* isolates, including paired colonizing and disease-causing multidrug-resistant isolates, was performed using Snippy v4.6 with the earliest isolate as the reference. For core genome SNP analysis, variants within repetitive regions (identified with nucmer from MUMmer) or prophage regions (PHASTER) were excluded. Maximum-likelihood phylogenetic trees were generated with IQ-TREE (GTR+G model, 1,000 bootstrap replicates). Pairwise SNP distances were visualized using the *pheatmap* package in R.

### Rifampin mutation frequency assays

Hypermutability was assessed by plating serial dilutions (∼10¹⁰ CFU/mL) on rifampin-containing (300 µg/mL) and drug-free MH agar.^52^ Mutation frequency was calculated as the ratio of colonies on rifampin plates to those on drug-free plates. Each isolate was tested in triplicate, and the mean ± SEM mutation frequency was reported.

### *In vitro* serial passage

Serial passaging was performed as described, with modifications.^53^ Briefly, experiments were carried out in 96-well plates with 2-fold serial dilutions of antibiotics in Mueller–Hinton (MH) broth. Ceftazidime and MEM were tested at concentrations ranging from 0.03 to 1,024 µg/mL with fixed 4 µg/mL avibactam or 8 µg/mL vaborbactam concentration, respectively. Each well was inoculated with ∼4×10⁶ cells, incubated aerobically at 35 °C for 18–24 h, and growth (OD₆₀₀ > 0.2) from the highest antibiotic concentration was diluted 1:100 into fresh medium. This process was repeated daily for 14 passages. MICs were determined every two passages by broth microdilution.

### Construction of *ΔmutH* mutant

A *mutH* deletion mutant was generated using a CRISPR/Cas9-mediated genome editing system.^26^ The plasmid pCasKP-apr was electroporated into strain D3 and selected on LB agar containing 30 µg/mL apramycin, followed by PCR confirmation. An sgRNA targeting *mutH* was designed using the SSC online server (http://crispr.dfci.harvard.edu/SSC/) and cloned into pSGKP-Rif, yielding pSGKP-mutH-N20. A 413-bp deletion repair template was amplified by two-step PCR using primers mutH-A/B/C/D. The repair DNA and pSGKP-mutH-N20 were co-transformed into L-arabinose–induced, pCasKP-apr–positive D3 cells. Cultures were plated on LB agar containing 30 µg/mL apramycin and 100 µg/mL rifampin, and candidate mutants were screened by PCR and confirmed by sequencing. To cure editing plasmids, mutants were grown in antibiotic-free LB broth and streaked on LB agar with 5% sucrose; colonies retaining the deletion but lacking both pCasKP-apr and pSGKP-RFP were confirmed by the inability to grow on antibiotic-containing plates.

### Allelic replacement of *mutH* with *mutH*^V76G^

*mutH*^V76G^ along with ∼500 bp of flanking upstream and downstream regions was amplified from isolate D5 DNA using primers KpMutH-complement-For and KpMutH-complement-Rev [Table 3]. The PCR product was cloned into the *Not*I site of the pKOV vector and verified by Sanger sequencing. The resulting recombinant plasmid, pKOV-*mutH*-V76G, was co-electroporated with pSGKP-mutH-N20 into the L-arabinose-induced *D3* isolate harboring the pCasKP-apr plasmid. Successful allelic replacement of native *mutH* locus with the *mutH*^V76G^ variant was confirmed by PCR and WGS. Plasmids were cured as described above.

**Table 3.**
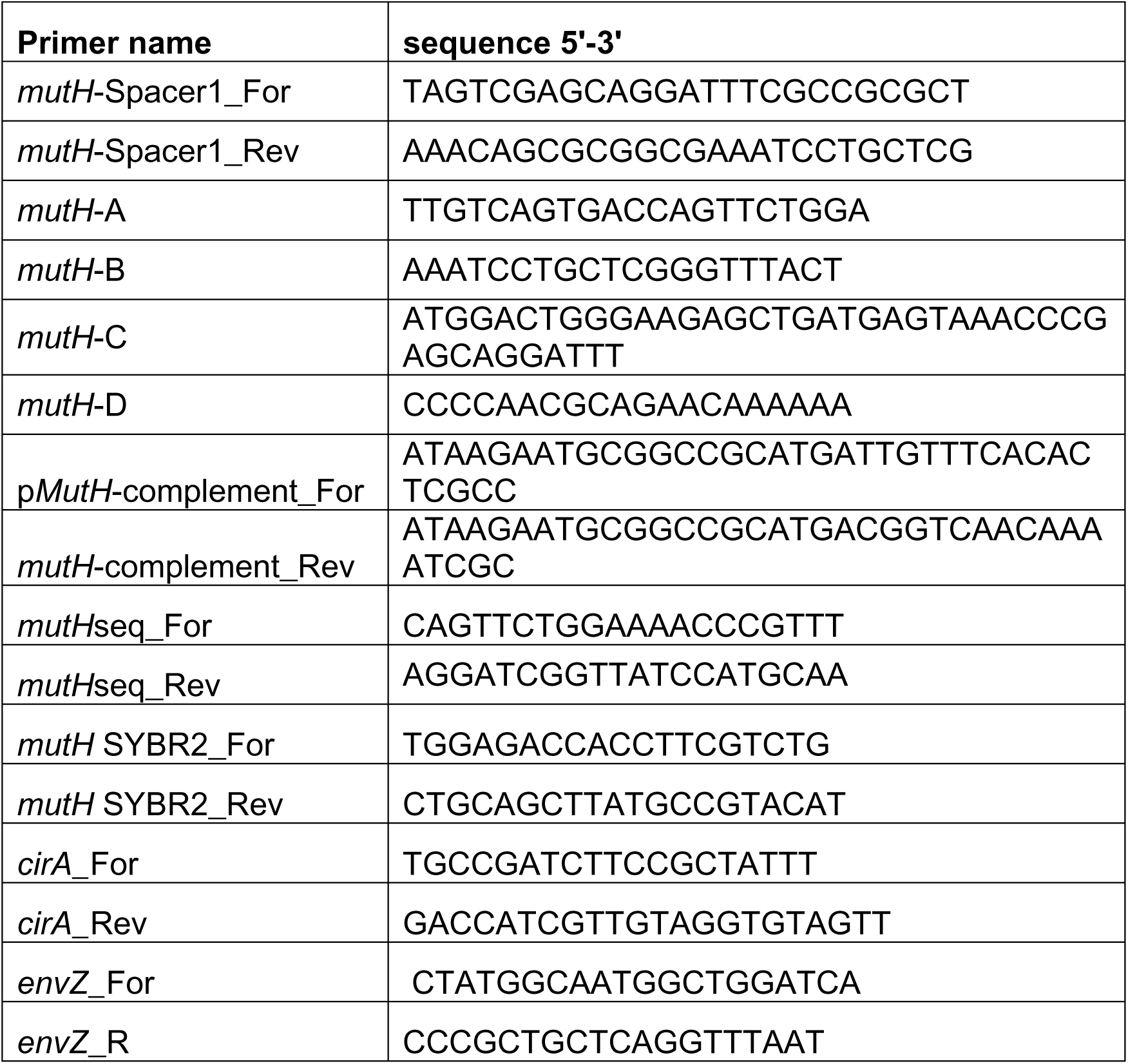
Primers used in this study.

### Conjugation assays

Conjugation experiments were performed using a broth mating protocol adapted from Huang et al.,^28^ with minor modifications. Donor and recipient strains were grown in Mueller–Hinton (MH) broth with appropriate antibiotics at 37°C to mid-exponential phase, washed three times with fresh MH, and adjusted to OD₆₀₀ ∼ 0.5. Equal volumes (0.5 mL) of donor and recipient cultures were mixed (1:1), pelleted, resuspended in 50 µL MH, and 25 µL was spotted onto 0.45 µm membrane filters on MH agar. Plates were incubated overnight at 37°C. After incubation, cells were recovered, serially diluted, and plated on CHROMagar with selective antibiotics. Controls included donor and recipient strains plated separately. *<u>Plasmid uptake (“recipient” assay)</u>:* Recipient isolate D3 (WT *mutH*) and Δ*mutH*, and *mutH*^V76G^ mutant strains were mated with *E. coli* XDR-74 carrying IncX3-*NDM-5* and selected with 50 mg/L kanamycin, and 16 mg/L ceftazidime with 4 mg/L avibactam. *<u>Plasmid donation (“donor” assay)</u>:* Donor strains D3::IncX3-*NDM-5*, Δ*mutH*::IncX3-*NDM-5*, and *mutH* V76G::IncX3-*NDM-5* were mated with *E. coli* J53::pCasApr (apramycin resistance) and selected with 100 mg/L apramycin, and 16 mg/L ceftazidime with 4 mg/L avibactam. Plates were incubated at 37°C for 18–24 h and colonies were enumerated, and the presence of *NDM* was confirmed by PCR. Conjugation frequency was calculated as the number of transconjugants per donor. All experiments were performed in triplicate.

### Mouse model of disseminated *K. pneumoniae* infection

Male and female ICR CD1 mice (20–25 g, Envigo) were used in equal number. Transient immunosuppression was induced via intraperitoneal cyclophosphamide (150 mg/kg 4 days prior and 100 mg/kg 1 day prior to infection) and subcutaneous cortisone (20 mg/kg 4 days prior and 1 day prior). Mice were intravenously inoculated via the lateral tail vein with 5×10⁴ (tissue burden studies) or 1×10^5^ (emergence of antifungal resistance studies) CFU of *K. pneumoniae*. For tissue burden tissue studies, mice were sacrificed on day 3, and liver, kidney and spleen were collected for colony enumeration. For emergence of resistance studies, the mice were treated with MVB (50 mg/kg intraperitoneally every 8 h) or CZA (37.5 mg/kg intraperitoneally every 8 h) starting at 4 hours post-infection. On day 3, mice were sacrificed, and liver, kidney, and spleen were homogenized and plated on LB agar with and without 4xMIC to screen for resistant isolates (CZA concentration: 1 µg/ml ceftazidime with 4 µg/ml avibactam; MVB concentration: 0.5 µg/ml MEM with 8 µg/ml vaborbactam). Tissue burdens were log_10_-transformed and expressed as mean ± SEM log_10_ CFU/gram of tissue. The difference between groups were determined by Wilcoxon test. *p-*value <0.05 was considered statistically significant.

### Statistical analysis

Data were analyzed using GraphPad Prism, v10.6.0 (GraphPad Software). When appropriate, data were log-transformed. Data were presented as mean ± SEM. Continuous variables were compared using Mann-Whitney U or Wilcoxon tests. For comparison of >2 groups, one-way ANOVA with Dunnett’s multiple comparisons test was used. Statistical significance was defined as *p-*value <0.05 (2-tailed).

## Ethical approval

All animal procedures were approved by the Institutional Animal Care and Use Committee (IACUC) at the University of Pittsburgh (Protocol #24014416) and were conducted in accordance with NIH guidelines for the care and use of laboratory animals. The study involving human subjects was approved by the Institutional Review Board (IRB #STUDY22040015), with a waiver of informed consent.

## Data availability

All genome sequences generated in this study have been deposited in the NCBI Sequence Read Archive (SRA) under BioProject accession number PRJNA1335250. Other data supporting the findings of this study are available from the corresponding author upon reasonable request.

## Acknowledgement

We acknowledge funding support from the National Institutes of Health (R21A1166847 awarded to M.H.N.), which made this work possible. We also extend our appreciation to Ryan Shields for his contribution to sample collection. The isolates included in this analysis were originally collected for a prior study (CREST), that was supported by the Antibacterial Resistance Leadership Group (to M. H. N., National Institute of Allergy and Infectious Diseases grant UM1AI104681)

## Author Contributions

S.C.: extracted strain DNA; conducted molecular biology experiments, SDS-PAGE, and other *in vitro* and mouse experiments; carried out whole-genome sequence (WGS) data analyses; interpreted data, drafted and revised manuscript; and formatted tables and figures. G.F. and H.B.: assisted SC in WGS data analyses, drafting, editing and revision of the manuscript; and in preparing tables and figures with genomic data. M.C.: conducted structural modeling and analysis that informed interpretation of the V76G substitution, reviewed and edited manuscript. AN: conducted experiments in conjunction with S.C, and contributed to manuscript preparation. LC: carried out whole-genome sequencing and accompanying data analyses, assisted with CRISPR-Cas9 gene disruption, conjugation assay and editing manuscript. M.H.N., C.J.C.: carried out study conception and design; oversaw experiments and data analyses, and redrafted, edited and revised the manuscript.

